# Selectivity filter mutation in Na_V_1.5 promotes ventricular tachycardia

**DOI:** 10.1101/2025.05.19.654350

**Authors:** Zoja Selimi, Mikhail Tarasov, Xiaolei Meng, Patrícia Dias, Bianca Moise, Rengasayee Veeraraghavan, Przemysław B. Radwański

**Affiliations:** The Frick Center for Heart Failure and Arrhythmia, Dorothy M. Davis Heart and Lung Research Institute, College of Medicine, The Ohio State University Wexner Medical Center, Columbus, OH, US; Division of Pharmaceutics and Pharmacology, College of Pharmacy, The Ohio State University, Columbus, OH, US; Department of Biomedical Engineering, College of Engineering, The Ohio State University, Columbus, OH, US; Department of Physiology and Cell Biology, College of Medicine, The Ohio State University, Columbus, OH, US

**Author notes:** **Corresponding author:** Przemysław B. Radwański, Pharm.D., Ph.D., Davis Heart and Lung Research Institute, The Ohio State University, 2255 Kenny Rd, Pelotonia Research Center, Columbus, OH 43210, Office.

**Keywords:** sodium channels, selectivity filter, Loss-of-Function, arrhythmia, Brugada syndrome

## Abstract

Loss-of-Function (LoF) mutations in the *SCN5A* gene, which encodes for the predominant cardiac Na_V_ isoform, Na_V_1.5 result in either deficiency in the channel expression or function. Impaired Na_V_1.5 expression and function underlie reduced peak Na^+^ current (I_Na_) and result in ventricular conduction velocity slowing, predisposing the heart to conduction block and ventricular arrhythmias clinically associated with Brugada syndrome (BrS). Recently, a missense mutation in Na_V_1.5 selectivity filter (DEKA motif), K1419E (DE**E**A) has been identified in patients with BrS. Despite early characterization of mutations in selectivity filter of other Na_V_ isoforms, little is known about the impact of DE**E**A on Na_V_1.5 function as well as on cardiac electrophysiology. Therefore, we generated a mouse heterozygous for Na_V_1.5 DE**E**A to characterize the mutation and investigate the outcome of this functionally deficient Na_V_1.5 variant on cardiac electrophysiology and arrhythmias. Heterologous expression system and isolated cardiomyocytes revealed lower current density and unchanged Na_V_1.5 expression in DE**E**A vs. wild type (DEKA). On the organ level, optical mapping revealed conduction velocity slowing in DE**E**A hearts, which was accentuated by flecainide resulting *in vivo* ventricular arrhythmias. Overall, to our knowledge, we provide the first mechanistic insight into the proarrhythmic consequences of a functionally deficient BrS mutation in Na_V_1.5.

**Condensed abstract:** Na_V_1.5 mutations have been associated with life-threatening arrhythmias. Recently, a selectivity filter mutation (K1419E-Na_V_1.5, DEKA→DE**E**A), has been linked to Brugada Syndrome (BrS). While DEKA mutations in other Na_V_ isoforms affected channel conductance, the impact of DE**E**A on Na_V_1.5 and arrhythmogenesis is unknown. Therefore, we generated mice heterozygous for Na_V_1.5-DE**E**A. Cardiomyocytes isolated from DE**E**A hearts exhibited substantial reduction in sodium current, ventricular conduction slowing and susceptibility to ventricular arrhythmias in vivo that were unmasked by flecainide. Together, DE**E**A murine model is the first to recapitulate a functional deficiency in Na_V_1.5, and thus offers insight into the proarrhythmic mechanism of BrS.

## Introduction

The cardiac sodium (Na^+^) channel Na_V_1.5 plays a critical role in mediating the inward Na^+^ current (I_Na_) that is responsible for the initiation of action potentials in the heart^1^. The permeability of Na_V_1.5 to Na^+^ is governed by key amino acid residues (DEKA) that compose the selectivity filter which line the pore of the channel^2–5^. Loss-of-function (LoF) mutations in *SCN5A*, the gene that encodes Na_V_1.5, have been associated with arrhythmias such as Brugada Syndrome (BrS)^6–8^ and a high risk of sudden cardiac death^9^. LoF mutations in *SCN5A* can result in a decreased Na_V_1.5 expression or produce a functionally defective Na_V_1.5 channels, both leading to reduced I_Na_. A decrease in I_Na_, in turn, compromises cellular excitability and slows electrical impulse propagation, which manifests as conduction velocity (CV)^6^ slowing. Collectively, these abnormalities in cardiac electrical impulse propagation create a substrate for reentrant arrhythmias^10,11^. A mutation affecting the Na_V_1.5 selectivity filter (DEKA) consisting of a substitution of a positively charged lysine residue at the position 1419 (K1419) with a negatively charged glutamic acid (DE**E**A) has been identified in BrS patients^12^. Evidence provided by studies in homologous variants in the selectivity filter of Na_V_1.2 and Na_V_1.4^3,4^ suggest that DE**E**A variant of Na_V_1.5 is expected to alter the permeability of the channel to Na^+^, thereby reducing overall I_Na_. One attempt to characterize the biophysical properties of DE**E**A Na_V_1.5 variant failed to yield detectable currents^13^. Furthermore, the electrophysiological consequences of this mutation on the organ level and *in vivo* have not been established. Therefore, we generated a BrS mouse model harboring the human homolog DE**E**A (K1421E). Contrarily to previous BrS mouse models which focused on the impact of haploinsufficiency or expression deficient variants that reduced I_Na_ only by 40-50%^14–18^, our murine model is the first model of a functionally deficient Na_V_1.5 channel variant. Importantly, DE**E**A cardiomyocytes evidenced a >70% reduction in I_Na_ compared to wild type controls (DEKA), while no alterations were detected in protein surface expression. Moreover, the functional Na_V_1.5 deficiency associated with DE**E**A variant contributed to ventricular CV slowing in murine hearts. Notably, CV slowing evidenced in DE**E**A hearts was further exacerbated by flecainide which promoted ventricular tachycardia (VT) *in vivo*. In short, our study provides a mechanistic insight into the proarrhythmic nature of a BrS-associated mutation in Na_V_1.5 selectivity filter (DE**E**A).

## Methods

### Cell culture and transfection

Chinese hamster ovary (CHO) - K1 cells (ATCC, Manassas, VA, USA) cells were cultured in F-12 medium supplemented with glutamine (Thermo Fisher Scientific), 10% heat-inactivated fetal bovine serum (Thermo Fisher Scientific) and 0.5% penicillin-streptomycin solution 10,000 UI/ml (Thermo Fisher Scientific) at 37°C in 5% CO_2_. CHO-K1 cells were transiently transfected in 24 well plates with 1 µg per well of pcDNA3.1(+)P2A-eGFP generated by GeneScript (Piscataway, NJ, USA) encoding for either human WT-Na_V_1.5 (DEKA; accession no.: NM_001099404.2) or human K1419E-Na_V_1.5 (DE**E**A; accession no.: NM_001099404.2:c.4255A>G) using Lipofectamine 3000 (SignaGen Laboratories). Green fluorescent protein (GFP) positive cells were used for patch-clamp experiments 48-72 hours post-transfections.

### Mouse model

Heterozygous mice harboring human Na_V_1.5 DE**E**A homolog (K1421E) were generated using the Neomycin (Neo) selection cassette for the conventional knock-in model by inGenious Targeting Laboratory Inc. Initially, targeted iTL BF1 (C57BL/6 FLP) embryonic stem cells were microinjected into Balb/c blastocysts. The resulting chimeras with a high percentage of black coat color were mated with C57BL/6N mice to generate germline Neo-deleted mice. The Neo-deletion was identified using the primer set PNDEL1/PNDEL2, which was confirmed by sequencing of the PCR product, thus, identifying a 450 bp band for Neo-deletion and an 83 bp band for the remaining FRT site. Next, to verify the presence of the FLP transgene, using the set of primers newFLP1/newFLP2. The intended point mutation (A→G) was verified through sequencing of the PCR product amplified by SQ1/RNEOGT primers. The heterozygous line was bred from a founder and maintained on a C57BL/6J (The Jackson Laboratory-Strain #:000664) background.

### Murine cardiomyocyte isolation

Ventricular cardiomyocytes were isolated from mice according to previously established protocols^19–21^. Mice were first anesthetized using 3-5% isoflurane in 100% oxygen (0.8-1.0 L/min), and once deep anesthesia was confirmed by the absence of the pedal reflex, mice were euthanized by cervical dislocation before the thoracotomy was performed. The heart was quickly removed and placed in cold Ca²⁺-free Tyrode’s solution (in mmol/L: 133.5 NaCl, 4 KCl, 1 MgCl₂, 10 glucose, and 10 HEPES, pH adjusted to 7.4 with NaOH). The aorta was then cannulated, and the heart was transferred to a Langendorff apparatus for perfusion with Ca^2+^-free Tyrode’s solution at 37°C to wash out the remaining blood. Next, the heart was perfused with Liberase TH (Roche). The ventricles were separated from the atria, minced in Tyrode’s solution containing 2% BSA, gently agitated to disperse the tissue, and filtered through a nylon mesh to isolate cardiomyocytes. These cells were resuspended in low-Ca^2+^ Tyrode’s solution (in mmol/L: 133.5 NaCl, 4 KCl, 1 MgCl₂, 0.1 CaCl₂, 10 glucose, and 10 HEPES, pH 7.4 adjusted with NaOH), and stored at room temperature for up to 5 hours.

### Whole-cell patch-clamp recordings

Whole-cell patch clamp recordings in CHO cells were acquired at room temperature using an Axopatch 200B amplifier and Digidata 1550B (Molecular Devices). The external solution to record inward Na^+^ currents (I_Na_) contained (in mmol/L): NaCl, 135; CsCl, 4; CaCl_2_, 1.8; MgCl_2_, 1.2, HEPES, 10; glucose, 10 with a pH=7.4 adjusted with NaOH, whereas the solution to assess Ca^2+^ permeability contained (in mmol/L): N-methyl-d-glutamine (NMDG), 120; CaCl_2_, 10; CsCl, 4; MgCl_2_, 1.2; HEPES, 10; glucose, 10 with pH=7.4 adjusted with HCl. The internal solution contained (in mmol/L) NaCl, 10; CsF, 130; EGTA, 10; HEPES, 10 at a pH=7.3 adjusted with CsOH. Whole-cell peak I_Na_ in isolated cardiomyocytes were acquired using a MultiClamp 700B amplifier and Digidata 1440A (Molecular Devices). For peak I_Na_, the bath solution was modified by reducing the NaCl to 10 mmol/L and increasing CsCl to 130 mmol/L. The composition of the internal solution was the same as above-described for CHO cells. Borosilicate pipettes (Sutter Instruments) with a resistance from 1.0-3.0 MΩ were pulled using a laser P2000 puller (Sutter Instruments). Whole-cell currents were low-pass filtered using a 5 kHz 4-pole Bessel filter and recorded at a sampling rate of 20 kHz. The liquid junction potential was corrected. Currents were recorded 5-10 minutes after achieving the whole-cell configuration. Steady-state activation and inactivation protocols (Figure 1C) were run to assess the kinetics of the currents. Recordings were analyzed using the Clampfit software (Molecular Devices) and custom-written MATLAB code (MathWorks, Natick, MA, USA). The activation and inactivation parameters for I_Na_ were fitted with a sigmoidal Boltzmann function 1/(1+exp((V-V_1/2_)/k)), where V is the membrane potential, V_1/2_ is the half activation voltage and k is the slope factor.

**Figure 1.**
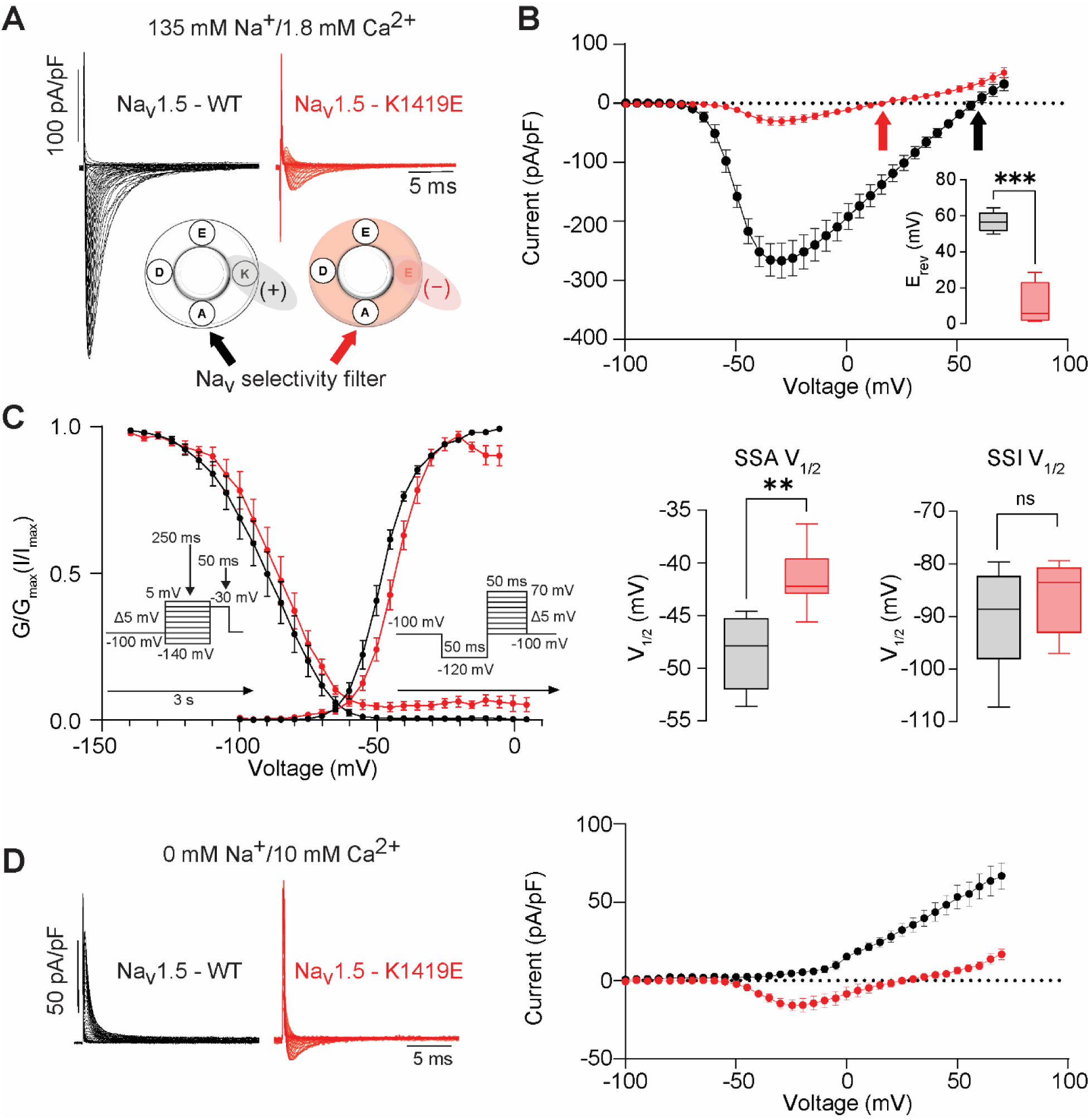
Na_V_1.5-K1419E (DE**E**A) exhibits reduced peak Na^+^ current (I_Na_) and altered ion selectivity. (A) Representative I_Na_ from wild type Na_V_1.5 (DEKA; black) and DE**E**A (red) and corresponding (B) I–V relationship, reversal potential (inset), (C) steady–state activation and inactivation (n = 8 and 7 for DEKA and DE**E**A, respectively). (D) Ca^2+^ permeability (n = 4 and 5 for DEKA and DEEA, respectively; *p≤0.05, *** p≤0.001).

### Fluorescent immunolabeling and confocal microscopy

Immunofluorescent labeling of 5 μm tissue sections was performed as previously described^20–22^. Briefly, the tissue sections were fixed using 2% PFA (MilliporeSigma) and then permeabilized for 15 minutes at room temperature using 0.2% Triton X100 (MilliporeSigma) followed with the blocking agent (1% bovine serum albumin, 0.1% triton in PBS) for 2 hours at room temperature. Next, the tissue sections were incubated overnight at 4°C with primary antibodies against N-cadherin (Ncad BD Biosciences 610920) and Na_V_1.5 (a previously validated custom rabbit polyclonal antibody^23^) channels. Samples were then washed 5×5 minutes in PBS and incubated with secondary antibodies (goat anti-rabbit Alexa Fluor 568 and goat anti-mouse Alexa Fluor 488 at 1:4,000; Thermo Fisher Scientific) for 2 hours at room temperature. Samples were again washed in PBS (5×5 minutes) and mounted in ProLong Gold (Thermo Fisher Scientific).

Confocal imaging was performed with a Nikon AX/R laser-scanning confocal microscope equipped with 4 solid-state lasers (405, 488, 560, and 647 nm, 30 mW each), a 63×/1.4NA oil-immersion objective, a spectral detection module comprised of 2 GaAsP detectors and 2 high-sensitivity photomultiplier tube detectors, a cutting-edge high-definition confocal scanner, and a high-speed piezo z-drive.

### Optical mapping

Optical mapping of Langendorff-perfused hearts was performed as described in previous studies^22,24^. Isolated murine hearts were perfused as Langendorff preparations with a solution containing (in mmol/L): 140 NaCl, 4.2 KCl, 1.8 CaCl_2_, 5.5 Dextrose, 1 MgCl_2_, 1.2 NaH_2_PO_4_ at a pH 7.4 corrected with NaOH. Temperature was maintained at 36°C at all times. Murine hearts were perfused with the voltage-sensitive fluorophore, di-4-ANEPPS (7.5 μmol/L, Biotium) for 15 minutes and blebbistatin (10 μmol/L, Sigma-Aldrich) was administered continuously for motion control. Hearts were paced with an AgCl_2_ electrode placed septally at the epicardium. Hearts were paced at basic cycle length of 120 ms. The anterior epicardium was mapped using a MiCAM03-N256 CMOS camera (SciMedia: 256×256 pixels, field of view 11.0×11.0 mm, 1.8-kHz frame rate). Conduction velocity (CV) was measured at baseline conditions and after challenging the hearts with flecainide 1 µM (Thermo Fisher Scientific). Conduction velocity (CV) was assessed in both the longitudinal (CV_L_) and transverse (CV_T_) directions as the average of local CVs along the long axis and the average of local CVs along the short axis of the cardiac tissue, respectively. Activation time was defined as the time at which the first derivative of the action potential reaches its maximum, as previously described^25,26^. Activation maps and phase maps were analyzed using BV Workbench Version 4.0.2.

### *In vivo* surface electrocardiograms (ECGs)

Electrocardiograms (ECGs) (PL3504 PowerLab 4/35, ADInstruments) were obtained from mice anesthetized with isoflurane (1%-3% isoflurane plus 100% oxygen, 0.8-1 L/min). After 10 minutes of baseline recording, an injection of flecainide (40 mg/kg, MilliporeSigma) was administered intraperitoneally (i.p), and the recording continued for an additional 40 minutes. The QRS complex duration was defined as the duration between the earliest moment of deviation from baseline up to the moment when the S-wave returned to the isoelectric line in lead I^22^. Ventricular tachycardia (VT) was defined as 3 or more consecutive premature beats ^20,22,27^. ECG recordings were analyzed using the LabChart 7.3 (ADInstruments).

### Statistical analysis

Statistical analyses were conducted using GraphPad Prism 10 (GraphPad Software). Data normality was assessed using the Shapiro-Wilk test, and the appropriate statistical tests were selected to compare the different groups. For comparing two independent datasets, unpaired, two-tailed Student’s t-test or Mann-Whitney U test was applied for normally and non-normally distributed data, respectively. Comparing more than two datasets, either a one-way ANOVA or Kruskal-Wallis test were performed for normally and non-normally distributed data, respectively. A p value lower than 0.05 was considered statistically significant. All data are presented as mean ± SEM or as box-and-whisker plots, where the box represents the first and third quartiles, the line within the box indicates the median, and the whiskers show the minimum and maximum values, unless otherwise stated. “n” represents the number of cells, and “N” denotes the number of mice.

## Results

### DEEA variant reduces I_Na_ and alters Na^+^ permeability of Na_V_1.5 in heterologous expression systems

A mutation in the Na_V_1.5 selectivity filter (DE**E**A) has been reported in a BrS patient^12^. However, the functional consequences of this mutation in the heart are unknown. Thus, prior to assessing the pro-arrhythmic potential of DE**E**A in the mouse heart, we examined how this mutation alters Na_V_1.5 properties, by transiently expressing the wild-type (DEKA) or K1419E variant (DE**E**A) channel in CHO cells (Figure 1) so that we could fully assess the Na^+^ current reduction. Consistent with previous findings in the Na_V_1.2 and Na_V1_.4 isoforms^3,4,28^, our results showed a drastic reduction in current density at all potentials in cells expressing the Na_V_1.5 DE**E**A variant compared to cells expressing Na_V_1.5 DEKA (WT) channels (-29.78±17.73 pA/pF vs. -263.02±73.08 pA/pF at -35 mV, p<0.0003) (Figure 1B). Furthermore, the reversal potential was left-shifted (12.24±10.43 vs. 56.90±4.77 mV, p<0.0003; Figure 1B inset), which suggests that DE**E**A channel lacks ion selectivity, being thus permeable to ions other than Na^+^. Indeed, DE**E**A evidenced permeability to Ca^2+^ when Na^+^ was removed from the pipette (−11.55±3.58 pA/pF vs. 2.65±2.04 pA/pF at -35 mV, p=0.0159; Figure 1D). Additionally, the activation kinetics was shifted to depolarizing potentials (-41.51±2.70 mV vs. -48.62±3.28 mV, p=0.0012), while steady-state inactivation was not affected by DE**E**A (Figure 1C).

### Reduction of I_Na_ in DEEA murine hearts

Although other Na_V_1.5 functional deficiencies have been extensively studied in heterologous expression systems^29,30^, studies in a physiological milieu are lacking. To uncover functional consequences of DE**E**A in mice, we generated a knock-in mouse with the murine homolog of DE**E**A, Scn5a-K1421E (Figure 2A). Consistent with observations in heterologous expression system, compared to DEKA (WT), DE**E**A cardiomyocytes exhibited reduced I_Na_ (−48.54±3.61 pA/pF vs. −10.467±2.77 pA/pF, p<0.0003 at -35 mV; Figure 2B), with a right-shift in activation kinetics (−51.81±4.09 mV vs. −43.18±2.61 mV, p=0.0219) with unaltered inactivation kinetics (Figure 2B). Despite the loss-of-function (LoF) phenotype evidenced in DE**E**A hearts, we did not observe changes in Na_V_1.5 expression (Figure 2C). Thus, our observations suggest that DE**E**A results in functional deficiency without observably altering channel expression.

**Figure 2.**
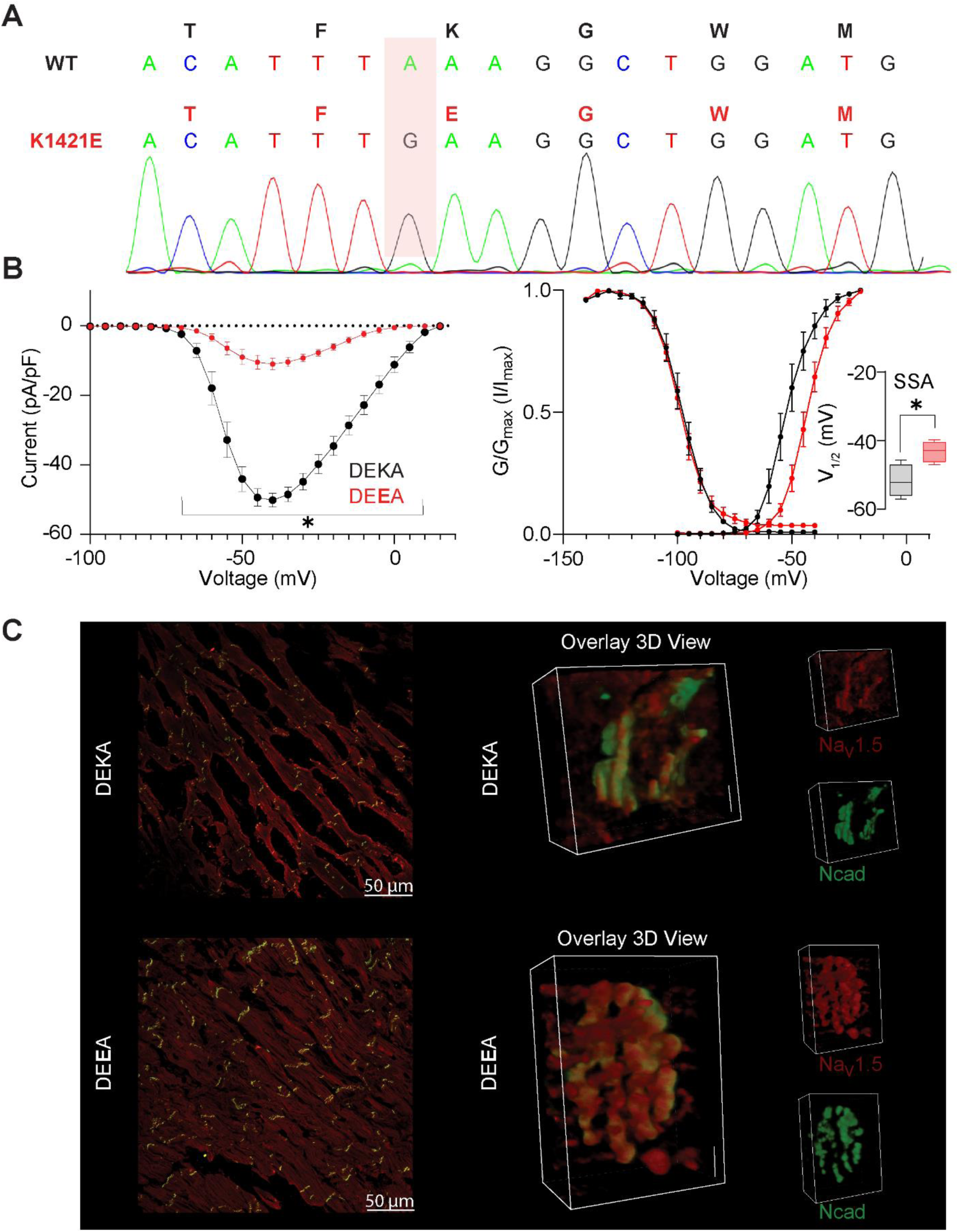
Murine DE**E**A homolog Na_V_1.5-K1421E exhibits reduced peak I_Na_. (A) Sequencing of *Scn5a* genomic PCR products with nucleotide substitution resulting in K1421E. (B) I–V relationship of wild type Na_V_1.5 (DEKA; black) and DE**E**A (red), (right) steady–state activation and inactivation (n = 4, N=2 for DEKA and n=4, N=2 for DE**E**A cardiomyocytes, respectively), *p≤0.05. (C) Distribution of Na_V_1.5 (red) and N-cadherin (Ncad; green) in WT (DEKA; top) and K1421E (DEEA; bottom) murine myocardium.

### DEEA mice exhibit cardiac conduction abnormalities and ventricular arrhythmias

To investigate the impact of DE**E**A-mediated functional deficiency on ventricular conduction, we performed optical mapping in DE**E**A and DEKA (WT) hearts. Our results showed a significant reduction in conduction velocity (CV) in DE**E**A vs. DEKA (WT) hearts both longitudinally along the muscle fiber (CV_L_; 47.29±0.29 cm/s vs. 60.21±1.53 cm/s, p=0.0053) as well as transversely (CV_T_; 31.65±4.39 cm/s vs. 40.8±1.80 cm/s, p=0.0146; Figure 3) without affecting the anisotropic ratio (1.49±0.09 vs. 1.48±0.03, p=0.8669). Next, to assess the conduction reserve, we perfused the hearts with 1 µM flecainide. As expected, flecainide slowed CV in both directions (Figure 3B); however, DE**E**A hearts evidenced a more pronounced reduction in CV_L_ (31.69±3.74% vs. 21.84±2.37%, p=0.0357) and CV_T_ (36.68±7.04% vs. 20.58±1.85%, p=0.0357) relative to DEKA hearts. Notably, the differences in CV observed between DE**E**A and DEKA hearts *ex vivo*, translated to QRS prolongation *in vivo* (10.56±0.99 ms vs. 8.49±0.51 ms, p=0.0462) (Figure 4A) which was further accentuated by flecainide challenge (40 mg/kg, i.p.; 23.05±3.72 ms vs. 17.24±2.89 ms, p<0.0003; Figure 4A). Importantly, the flecainide challenge promoted ventricular tachycardias (VT) in 23.52% (4 of 17 mice) of the DE**E**A mice, while no VTs were observed in DEKA (WT) mice (Figure 4B). Thus, to our knowledge, this is the first report of a functionally deficient Na_V_1.5 variant murine model that recapitulates BrS-associated arrhythmia phenotype.

**Figure 3.**
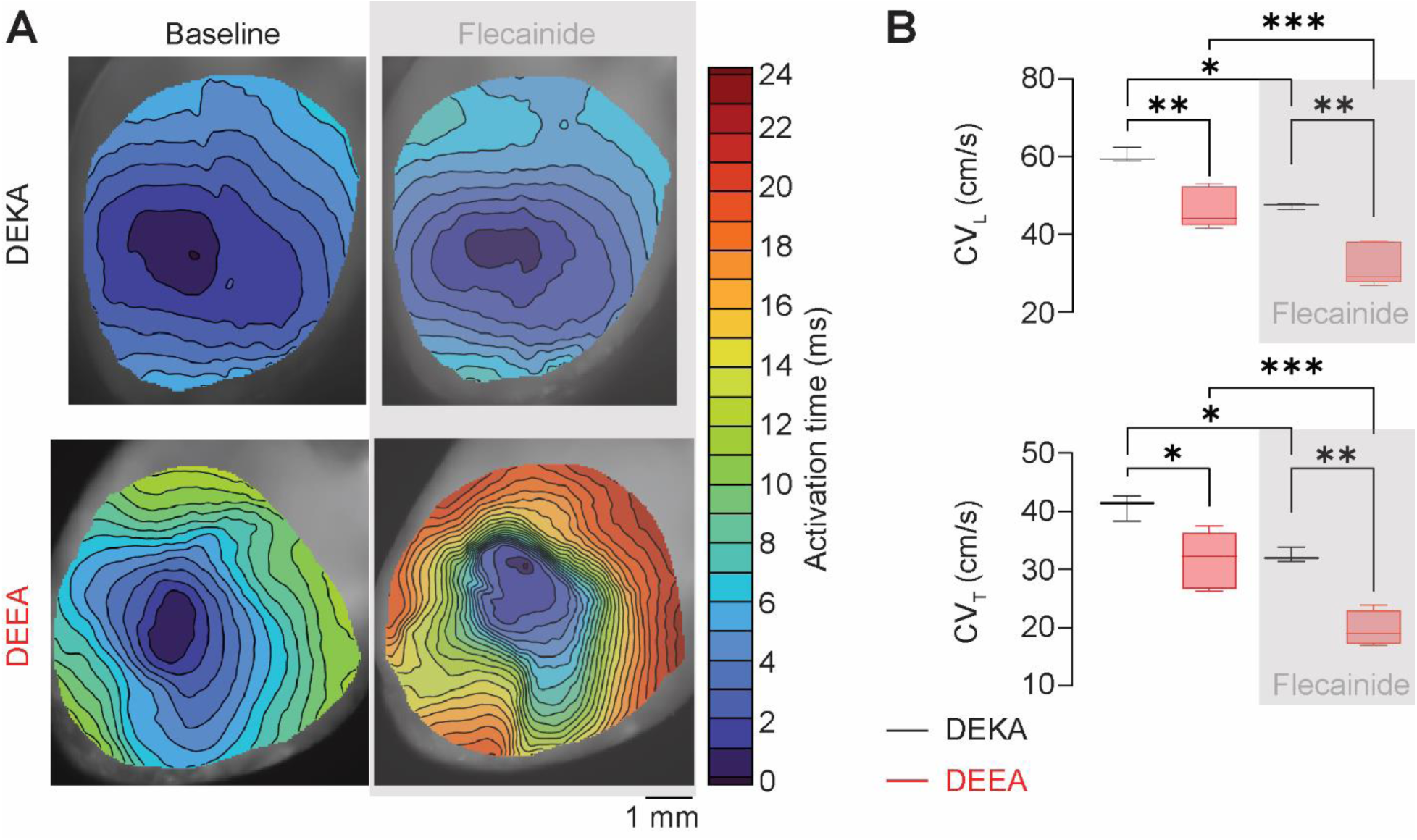
DE**E**A hearts exhibit a reduction in conduction velocity compared to DEKA hearts, which is further exacerbated by flecainide. (A) Representative isochrone activation maps from DEKA and DE**E**A hearts recorded during baseline and after flecainide (1 µM). (B) Conduction velocities in longitudinal (CV_L_) and transverse (CV_T_) directions, before and after the flecainide challenge (grey). (N=3 for DEKA and N=5 for DE**E**A) *p≤0.05, **p≤0.01, ***p≤0.001.

**Figure 4.**
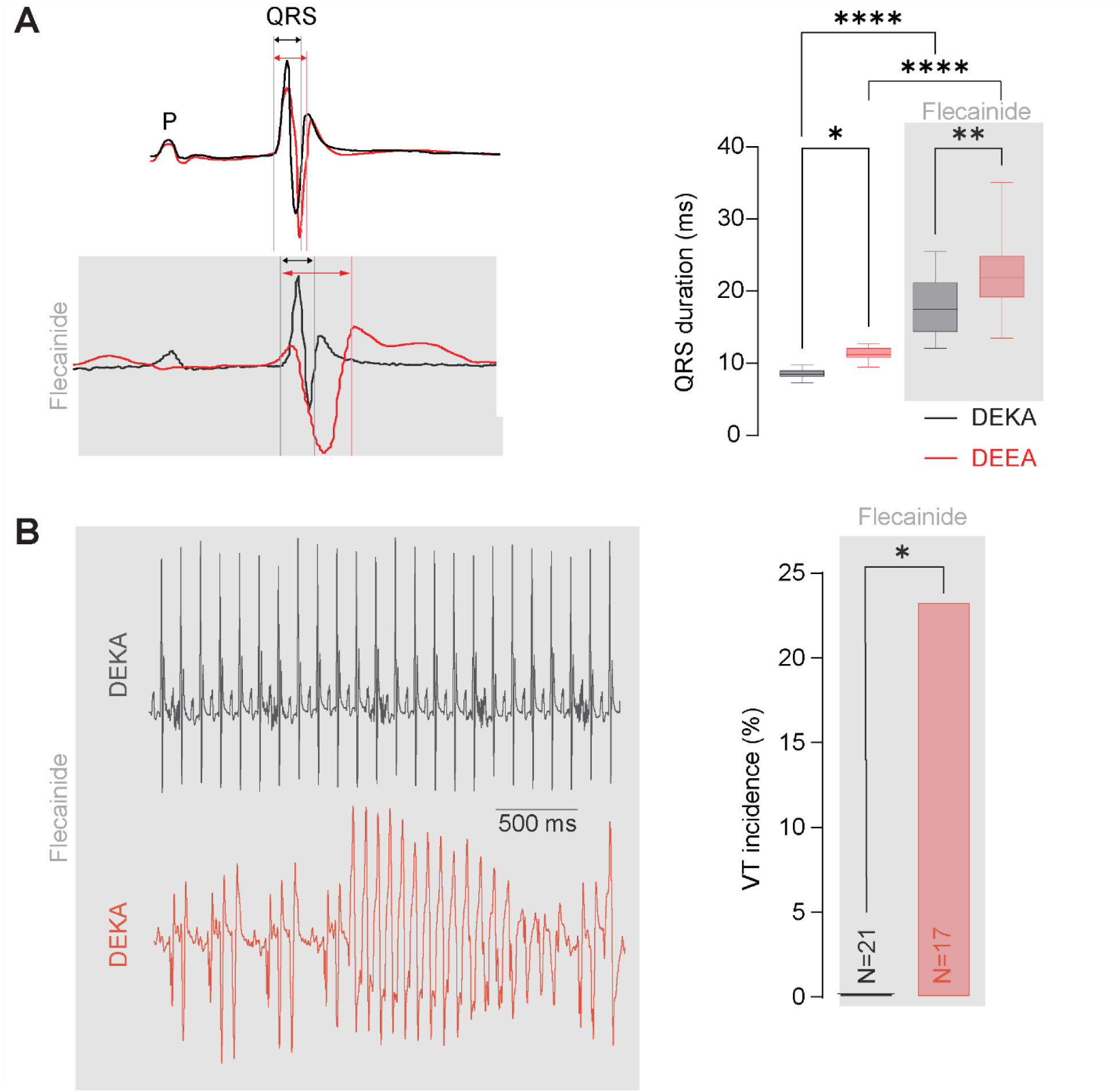
Flecainide unmasks VT in DE**E**A mice. (A) QRS interval is longer in DE**E**A (red) compared to DEKA (black) and it is prolonged after flecainide (40 mg/kg, i.p.) (N=21 for DEKA and N=17 for DE**E**A). (B) Flecainide unmasks VT in DE**E**A mice, N presented in the figure *p≤0.05, **p≤0.01, ***p≤0.001.

## Discussion

Loss-of-function mutations in *SCN5A* have been associated with BrS. These mutations lead to a reduced Na^+^ current density (I_Na_), either due to deficiency in surface expression or functional deficiency^6^. Recently, a BrS patient with a point mutation (K1419E) affecting the Na_V_1.5 selectivity filter (DE**E**A) has been identified^12^. Despite in-depth studies that helped develop a better understanding of the biophysical consequences of mutations in the selectivity filter in other Na_V_ isoforms^3,4,28^, the impact of DE**E**A on Na_V_1.5 cardiac isoform and arrhythmogenic consequences remain elusive. Therefore, we sought to uncover the functional consequences of DE**E**A mutation with regard to Na_V_1.5 channel behavior along with its impact on electrical impulse propagation and proarrhythmic potential in the myocardium. To that end, we generated a mouse model harboring DE**E**A, which recapitulates reduced I_Na_ observed in heterologous systems. Compromised Na_V_1.5 function promotes slowed cardiac conduction that is exacerbated by Na^+^ channel blockade with flecainide which unmasks VTs *in vivo*. To our knowledge, this is the first description of a murine model with a functionally deficient Na_V_1.5 variant that phenocopies BrS-associated VT.

Previous studies in *Scn5a* expression deficiency murine models have provided valuable insights into the pathophysiology of BrS. The first BrS mouse model to be generated was *Scn5a* haploinsufficient^14^, which resulted in 50% reduction in I_Na_, reduced CV and VTs^14^. Similarly, subsequent models, such as the 1798insD^17^, D1275N^18^ and G1746R^16^ also evidenced I_Na_ reduction by 40-50%. While 1798insD model exhibited a Long QT phenotype and reduced surface expression^31^, no VTs were reported upon flecainide challenge in mice on the FVB/N background^17^. Conversely, 1798insD mice on 129P2 background exhibited arrhythmias at rest, which were ascribed to increased late sodium current (I_NaL_) in that strain^32^. On the other hand, mice harboring the D1275N and G1746R expression deficient mutations were positive for VT phenotype^16,18^. However, the impact of DE**E**A functional deficiency had not yet been demonstrated in a complex milieu of a cardiomyocytes. Importantly, it is also not clear whether I_Na_ would be sufficiently impacted in a DE**E**A cardiomyocyte and whether this reduction in I_Na_ would be sufficient to phenocopy BrS-associated VT. To address this, we generated a murine model harboring a mutation in Na_V_1.5 selectivity filter (DE**E**A), which resulted in a functional defect accompanied by an >70% reduction in peak I_Na_. Our model highlights the robustness of the excitability reserve present in the mammalian heart, where a substantial reduction in I_Na_ is required in order to unmask the BrS-associated VT phenotype.

## Conclusion

In summary, the DE**E**A mutation in the Na_V_1.5 selectivity filter results in functional deficiency and substantial I_Na_ reduction. DE**E**A mice evidence compromised conduction velocity *ex vivo* and increased arrhythmic risk *in vivo* that is unmasked by sodium channel blockade. These findings underscore the importance of a large excitability reserve, which, when compromised, can result in ventricular arrhythmias. It is noteworthy that *Scn5a* haploinsufficiency and expression-deficient Na_V_1.5 mutations can be rescued by increasing surface expression^16^. Future studies should focus on development of therapies that rescues excitability reserve in functionally deficient Na_V_1.5 variants. Lastly, the DE**E**A mouse model described herein is a valuable platform for further exploration of *SCN5A*-related disorders and development of potential therapeutic approaches.

This work was supported by National Institutes of Health grants R01NS121234 and R01HL155378 (to Dr Radwański) and R01HL148736 (to Dr Veeraraghavan). The authors have reported that they have no relationships relevant to the contents of this paper to disclose.

## Acknowledgements

The authors thank Joseph Waterman for maintaining the mouse colony used in this study and Madison Ammon for heart extraction and preservation for the immunofluorescence studies.

## References

1. Li RA, Tomaselli GF, Marbán E. Chapter 1 - Sodium Channels. In: Zipes DP, Jalife J, editors. Cardiac Electrophysiology (Fourth Edition). W.B. Saunders, 2004:1–9.

2. Favre I, Moczydlowski E, Schild L. On the structural basis for ionic selectivity among Na+, K+, and Ca2+ in the voltage-gated sodium channel. Biophys J. 1996;71:3110–3125.

3. Heinemann SH, Terlau H, Stühmer W, Imoto K, Numa S. Calcium channel characteristics conferred on the sodium channel by single mutations. Nature. 1992;356:441–443.

4. Schlief T, Schönherr R, Imoto K, Heinemann SH. Pore properties of rat brain II sodium channels mutated in the selectivity filter domain. Eur Biophys J. 1996;25:75–91.

5. Sun YM, Favre I, Schild L, Moczydlowski E. On the structural basis for size-selective permeation of organic cations through the voltage-gated sodium channel. Effect of alanine mutations at the DEKA locus on selectivity, inhibition by Ca2+ and H+, and molecular sieving. J Gen Physiol. 1997;110:693–715.

6. Wilde AAM, Amin AS. Clinical Spectrum of SCN5A Mutations: Long QT Syndrome, Brugada Syndrome, and Cardiomyopathy. JACC Clin Electrophysiol. 2018;4:569–579.

7. Schulze-Bahr E, Eckardt L, Breithardt G, et al. Sodium channel gene (SCN5A) mutations in 44 index patients with Brugada syndrome: different incidences in familial and sporadic disease. Hum Mutat. 2003;21:651–652.

8. Tan HL, Bezzina CR, Smits JPP, Verkerk AO, Wilde AAM. Genetic control of sodium channel function. Cardiovasc Res. 2003;57:961–973.

9. Sarquella-Brugada G, Campuzano O, Arbelo E, Brugada J, Brugada R. Brugada syndrome: clinical and genetic findings. Genet Med. 2016;18:3–12.

10. Kléber AG, Rudy Y. Basic mechanisms of cardiac impulse propagation and associated arrhythmias. Physiol Rev. 2004;84:431–488.

11. Lieve KV, Wilde AA. Inherited ion channel diseases: a brief review. EP Europace. 2016;17:ii1–ii6.

12. Kapplinger JD, Tester DJ, Alders M, et al. An international compendium of mutations in the SCN5A-encoded cardiac sodium channel in patients referred for Brugada syndrome genetic testing. Heart Rhythm. 2010;7:33–46.

13. Mikhailova VB, Karpushev AV, Vavilova VD, et al. Functional Analysis of SCN5A Genetic Variants Associated with Brugada Syndrome. Cardiology. 2022;147:35–46.

14. Papadatos GA, Wallerstein PMR, Head CEG, et al. Slowed conduction and ventricular tachycardia after targeted disruption of the cardiac sodium channel gene Scn5a. PNAS. 2002;99:6210–6215.

15. Leoni A-L, Gavillet B, Rougier J-S, et al. Variable Na(v)1.5 protein expression from the wild-type allele correlates with the penetrance of cardiac conduction disease in the Scn5a(+/-) mouse model. PLoS One. 2010;5:e9298.

16. Yu G, Chakrabarti S, Tischenko M, et al. Gene therapy targeting protein trafficking regulator MOG1 in mouse models of Brugada syndrome, arrhythmias, and mild cardiomyopathy. Sci Transl Med. 2022;14:eabf3136.

17. Remme CA, Verkerk AO, Nuyens D, et al. Overlap syndrome of cardiac sodium channel disease in mice carrying the equivalent mutation of human SCN5A-1795insD. Circulation. 2006;114:2584–2594.

18. Watanabe H, Yang T, Stroud DM, et al. Striking In vivo phenotype of a disease-associated human SCN5A mutation producing minimal changes in vitro. Circulation. 2011;124:1001–1011.

19. Munger MA, Olğar Y, Koleske ML, et al. Tetrodotoxin-Sensitive Neuronal-Type Na+ Channels: A Novel and Druggable Target for Prevention of Atrial Fibrillation. J Am Heart Assoc. 2020:e015119.

20. Tarasov M, Struckman HL, Olgar Y, et al. NaV1.6 dysregulation within myocardial T-tubules by D96V calmodulin enhances proarrhythmic sodium and calcium mishandling. J Clin Invest. 2023;133:e152071.

21. King DR, Demirtas M, Tarasov M, et al. Cardiac-Specific Deletion of Scn8a Mitigates Dravet Syndrome-Associated Sudden Death in Adults. JACC: Clinical Electrophysiology. 2024;10:829–842.

22. Dias P, Meng X, Selimi Z, Struckman H, Veeraraghavan R, Radwański PB. Lamotrigine promotes reentrant ventricular tachycardia in murine hearts. Epilepsia. 2025. Published onlineJanuary 30, 2025. 10.1111/epi.18295.

23. Veeraraghavan R, Hoeker GS, Alvarez-Laviada A, et al. The adhesion function of the sodium channel beta subunit (β1) contributes to cardiac action potential propagation. Elife. 2018;7.

24. Struckman HL, Moise N, King DR, et al. Unraveling Impacts of Chamber-Specific Differences in Intercalated Disc Ultrastructure and Molecular Organization on Cardiac Conduction. JACC Clin Electrophysiol. 2023;9:2425–2443.

25. Radwański PB, Veeraraghavan R, Poelzing S. Cytosolic calcium accumulation and delayed repolarization associated with ventricular arrhythmias in a guinea pig model of Andersen-Tawil syndrome. Heart Rhythm. 2010;7:1428–1435.e1.

26. Radwański PB, Poelzing S. NCX is an important determinant for premature ventricular activity in a drug-induced model of Andersen-Tawil syndrome. Cardiovasc Res. 2011;92:57–66.

27. Koleske M, Bonilla I, Thomas J, et al. Tetrodotoxin-sensitive Navs contribute to early and delayed afterdepolarizations in long QT arrhythmia models. J Gen Physiol. 2018;150:991–1002.

28. Echevarria-Cooper DM, Hawkins NA, Misra SN, et al. Cellular and behavioral effects of altered NaV1.2 sodium channel ion permeability in Scn2aK1422E mice. Hum Mol Genet. 2022;31:2964–2988.

29. Kyndt F, Probst V, Potet F, et al. Novel SCN5A mutation leading either to isolated cardiac conduction defect or Brugada syndrome in a large French family. Circulation. 2001;104:3081–3086.

30. Frosio A, Micaglio E, Polsinelli I, et al. Unravelling Novel SCN5A Mutations Linked to Brugada Syndrome: Functional, Structural, and Genetic Insights. Int J Mol Sci. 2023;24:15089.

31. Stein M, van Veen TAB, Remme CA, et al. Combined reduction of intercellular coupling and membrane excitability differentially affects transverse and longitudinal cardiac conduction. Cardiovasc Res. 2009;83:52–60.

32. Rivaud MR, Baartscheer A, Verkerk AO, et al. Enhanced late sodium current underlies pro-arrhythmic intracellular sodium and calcium dysregulation in murine sodium channelopathy. Int J Cardiol. 2018;263:54–62.

